# MosaicLev: Modified Levenshtein distance for mobile element-aware genome comparison

**DOI:** 10.64898/2026.02.03.703520

**Authors:** Harry Stoltz, Thomas E. Kuhlman

**Author notes:** **Supplementary information** Supplementary data, analysis scripts, and pairwise distance matrices are available at the GitHub repository.

## Abstract

**Motivation:** Somatic genetic mosaicism arises when genomes diverge across cells during development, in part due to the activity of transposons (cut–paste) and retrotransposons (copy–paste). Standard sequence comparison methods are not motif-aware, penalizing mobile element insertions based on length rather than recognizing them as single biological events.

**Results:** We introduce a modified Levenshtein distance (mlev) that discounts mobile element insertions via a tunable parameter (*m* ∈ [0, 1]). Validation on 67 Cluster G1 mycobacteriophage genomes demonstrates bidirectional discrimination between Mycobacteriophage Mobile Element 1 (MPME1) and MPME2 elements: using MPME1 as target yields ∼49% discount for MPME1 carriers but only ∼9% for MPME2 carriers, while using MPME2 reverses this pattern. This approach classifies 35 MPME1 carriers and 11 MPME2 carriers, and identifies 14 phages showing low discount with both motifs, consistent with absence of both elements.

**Availability and Implementation:** Python implementation with Numba JIT compilation freely available at https://doi.org/10.5281/zenodo.18452982.

## Introduction

In this paper, we will provide a mathematical exploration into genetic mosaicism. **Genetic Mosaicism** is a condition (see (Gill et al., 1995; Biesecker and Spinner, 2013)) where two or more genetically distinct cell populations exist within a single organism. This can occur due to the accumulation of stochastic mutations during development or throughout the organism’s life. We will focus on mutations due to the activity of Transposons (Muñoz-López and García-Pérez, 2010) and Retrotransposons (Finnegan, 2012). Transposons, also called “jumping genes,” are DNA sequences which can move around the genome via a cut-and-paste mechanism. Retrotransposons are a kind of transposon which uses a copy-and-paste mechanism to propagate within a genome. As these mobile elements move within a genome, they generate a variety of mutations. LINE-1 is one of the most common retrotransposons in humans, comprising ∼17% of the human genome (Lander et al., 2001; Cordaux and Batzer, 2009), and has been seen to be upregulated in brain development (see (Muotri et al., 2005)).

A rigorous mathematical study of **genetic mosaicism** may address the following questions:

1. How can we formally represent the cut–paste and copy–paste mechanisms of transposons and retrotransposons?
2. What are the formal language properties of genomes generated by such mechanisms?
3. How should we quantify the “distance” between two genomes when some mutations (e.g., mobile element insertions) represent single biological events rather than many independent changes?
4. Can such a distance measure detect biologically meaningful signals in real genomic data?

Standard sequence comparison methods (Durbin et al., 1998) treat all mutations uniformly: a 400-base insertion costs 400 times as much as a single substitution, regardless of whether that insertion is a mobile element that inserted in a single biological event. Existing approaches cannot fill this gap: affine gap penalties discount multi-base indels but cannot distinguish insertion sequences; mobile element detection tools identify insertions but do not produce pairwise distances or detect positional polymorphism. This paper addresses these questions by representing mosaicism as a finite rewrite system and introducing a modified edit distance with a parameter *m* ∈ [0, 1] controlling how strongly block events are discounted.

### Contributions

- We formalize transposon and retrotransposon mechanisms as a “mosaicism grammar” and analyze the induced language.
- we define a modified edit distance mlev with tunable block-event discounting and provide a dynamic-programming recurrence.
- we validate on 67 mycobacteriophage genomes (2,211 pairs), confirming theoretical predictions and detecting positional polymorphism.
- we demonstrate bidirectional discrimination between MPME1 and MPME2: running mlev with MPME2 as target classifies 11 MPME2 carriers and identifies 14 phages with low discount against both motifs, consistent with absence of both elements.

### Related work

Affine gap penalties (Gotoh, 1982) discount multi-base indels but treat all gaps equally regardless of sequence content. Block edit distances (Lopresti and Tomkins, 1997; Gańczorz et al., 2018) allow substring operations but are NP-hard in general and not motif-specific. Mobile element detection tools (MELT (Gardner et al., 2017), RetroSeq (Keane et al., 2013)) identify insertions from sequencing reads but do not produce pairwise distances. The mlev framework fills this gap by discounting only user-specified motif sequences while preserving full cost for other indels, enabling both quantitative distance computation and target-specific mobile element detection.

## Methods

### Mosaicism Grammar

We model mobile element activity as a string rewrite system (Book and Otto, 1993) over the DNA alphabet Σ = {*A, T, C, G*}. Transposons, also called “jumping genes,” are DNA sequences that move around the genome via a cut-and-paste mechanism: the element excises from one location and reinserts at another (Wicker et al., 2007). Retrotransposons use a copy-and-paste mechanism, propagating via an RNA intermediate that leaves the source copy intact while inserting a duplicate elsewhere. This biological asymmetry—transposons relocate while retrotransposons duplicate—directly motivates the cost structure in our distance metric: transposon sequences receive discounted costs for both insertion and deletion (reflecting coordinated cut-paste), while retrotransposon sequences receive discounted insertion but full per-base deletion cost (since removing an inserted copy is not a single biological event).

#### Truncated insertions

Retrotransposon insertion frequently produces 5’-truncated copies, thought to result from incomplete reverse transcription (Szak et al., 2002). We model this by allowing insertion of any suffix of the element sequence. For a retrotransposon string *R* = (*R*_1_)(*R*_2_) · · · (*R*_|*R*|_), truncated insertion can produce any suffix (*R*_*k*_) · · · (*R*_|*R*|_) for 1 ≤ *k* ≤ |*R*|. This motivates the suffix-based cost structure in our modified distance metric (Section 2.2).

#### Rewrite system formulation

We define the *expansion w*^*′*^ of a genome *w* = *a*_1_*a*_2_ · · · *a*_*n*_ ∈ Σ^∗^ to be the string 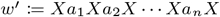, where *X* is a nonterminal symbol. Similarly, we write *T* ^*′*^ and *R*^*′*^ for the expanded forms of *T* and *R*. In the rule *T* ^*′*^ → *XIX*, the nonterminal *I* is linked with *T* ^*′*^; the swap rules allow us to shift *I* through the genome to the desired insertion point.

With these rules, we have the mosaicism grammar *G*. We write all the rules below for reference:

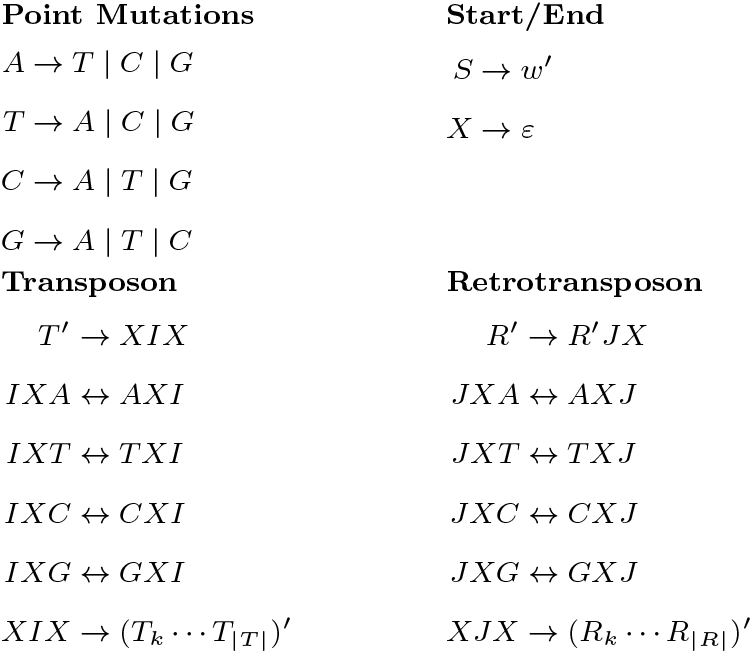

The final rules allow truncated insertion: any suffix (*T*_*k*_) · · · (*T*_|*T* |_) or (*R*_*k*_) · · · (*R*_|*R*|_) for 1 ≤ *k* ≤ |*T* | or |*R*|. The swap rules *IXA* ↔ *AXI*, etc., allow us to shift *I* through the genome to the desired insertion point; analogous rules hold for *J*.

#### Connection to the distance metric

The grammar’s suffix-based insertion rules directly motivate the cost structure of mlev (Section 2.2): the chunk sets ℳ_*T*_ and ℳ_*R*_ contain exactly those suffixes that can arise from truncated transposon and retrotransposon insertions in the grammar. The dynamic programming recurrence computes minimum-cost edit sequences that correspond to biologically plausible mutation pathways—sequences of point mutations and block-level mobile element events as formalized by the grammar rules above.

### Modified Levenshtein Distance

The classical Levenshtein distance function lev(*a, b*) (Levenshtein, 1966) is the minimum number of single-character insertions, deletions, or substitutions required to transform *a* into *b*. However, this unmodified Levenshtein function overemphasizes genomic effort with regards to retrotransposon and transposon cut-copy-paste operations. We want to treat insertions or deletions of mobile element strings as less costly than a series of single-character operations, but more than a sole single-character operation.

So, it makes sense to assign a cost that scales between 1 and the block length. We introduce a discount parameter *m* ∈ [0, 1] and define the cost of a block edit of length *L* by

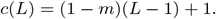

Thus *c*(1) = 1, *c*(*L*) = *L* at *m* = 0 (recovering classical Levenshtein), and *c*(*L*) = 1 at *m* = 1 (a whole block acts like a single edit).

Setup (chunks and costs)

We fix forward-oriented strings (Retrotransposons and Transposons)

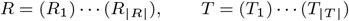

and a global (mosaicism) parameter *m* ∈ [0, 1] controlling the discount.

We then use the following “chunk sets:”

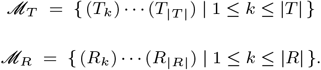

These are the right-end suffixes of the strings *R* and *T*.

#### Allowed primitive edits

Given words over Σ = {*A, T, C, G*}, we allow:

1. Single-base insertion, deletion, substitution (cost: 1).
2. *Block insertion* (transposons and retrotransposons): insert any chunk *M* ∈ ℳ_*T*_ ∪ ℳ_*R*_ (cost: *c*(|*M* |)).
3. *Block deletion (transposon only)*: delete any chunk *M* ∈ ℳ_*T*_ (cost *c*(|*M* |)).

We write cost(*e*) for the cost of a primitive edit *e*: single-base edits have cost 1, and any block insertion/deletion of a chunk *M* has cost *c*(|*M* |).

*Note: there is* no *discounted deletion for* ℳ_*R*_ *(retrotransposon chunks). Biologically, retrotransposons use a copy–paste mechanism, so the source copy remains in the genome; removing an inserted copy is not a single coordinated excision event like transposon cut–paste, and is therefore charged per base*.

### Definition

For *a, b* ∈ Σ^∗^, let ℰ (*a, b*) be the set of finite edit sequences using the primitives above. The modified distance is

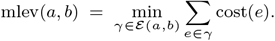

The distance mlev satisfies standard metric properties (non-negativity, definiteness, triangle inequality) but is asymmetric when *m >* 0 due to the transposon/retrotransposon distinction described above. Throughout this paper, we report the symmetric version 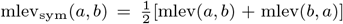 to ensure comparability across all pairwise analyses.

### Dynamic-programming formulation of the modified Levenshtein

Write *a* = *a*_1_ · · · *a*_*p*_ and *b* = *b*_1_ · · · *b*_*q*_.

Let *D*[*i, j*] be the optimal cost to transform *a*[1..*i*] to *b*[1..*j*], 0 ≤ *i* ≤ *p*, 0 ≤ *j* ≤ *q*, with *D*[0, 0] = 0.

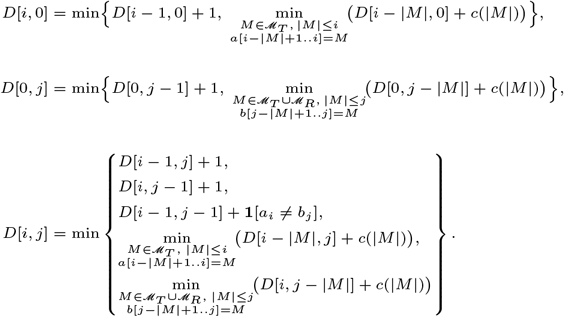

The value mlev(*a, b*) is *D*[*p, q*]. The DP runs in *O*(*pq*) time with precomputed suffix matches (Wagner and Fischer, 1974; Gusfield, 1997; Needleman and Wunsch, 1970; Navarro, 2001; Smith and Waterman, 1981). The implementation is available at https://doi.org/10.5281/zenodo.18452982, validated by property-based tests and parameter sweeps; mlev ≤ lev held in all cases.

#### Worked example

Let *T* = `TTAA` be a transposon motif, *R* = `ATG` a retrotransposon motif, and *m* = 0.5. Then *c*(3) = 1 + (0.5)(2) = 2 and *c*(4) = 1 + (0.5)(3) = 2.5. Consider:

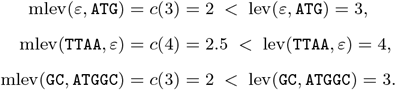

In the third case, the optimal edit sequence inserts the retrotransposon suffix `ATG` at cost *c*(3) = 2, rather than three single-character insertions at cost 3. At *m* = 1, all three block operations would cost exactly 1, treating each mobile element event as a single biological operation regardless of length.

### Dataset

We validated mlev on mycobacteriophage genomes containing a well-characterized mobile genetic element, providing biological ground truth: we know *a priori* which genomes contain the element, allowing verification that mlev correctly discounts mobile element insertions.

Mycobacteriophages are viruses that infect mycobacteria (Hatfull, 2010b). Cluster G phages (Pope et al., 2015) are closely related temperate phages with genome sizes of approximately 41–42 kb; some contain a mobile element called MPME1 (Mycobacteriophage Mobile Element 1, 439 bp), while others lack it entirely (Sampson et al., 2009). A related element MPME2 (440 bp) also exists, sharing ∼78% sequence identity with MPME1 (Sampson et al., 2009; Hatfull, 2010a). Phages lacking MPME1 have genomes of ∼41,441 bp, while those containing it have genomes of ∼41,901 bp (a difference of ∼445–460 bp reflecting the element plus flanking sequence). For computational analysis, we extracted a 434 bp core sequence from the BPs genome as our MPME1 motif (see Data Availability for coordinates).

We analyzed all 67 Cluster G1 mycobacteriophages available in PhagesDB (Russell and Hatfull, 2017) as of January 2025, representing the complete G1 subcluster. Cluster G comprises 82 phages across five subclusters (G1–G5) as of January 2025 (Russell and Hatfull, 2017); we focused on G1 because it is the largest and contains the well-characterized MPME1 and MPME2 elements. This yields 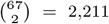 pairwise comparisons with no selection bias. Based on genome size, seven phages with short genomes (∼41,077– 41,456 bp) are inferred to lack mobile elements, while 60 phages with longer genomes (∼41,650–42,400 bp) are inferred to contain MPME1 or a related element (our mlev analysis below distinguishes MPME1 from MPME2 carriers and identifies phages lacking both elements):

This yields three comparison categories: 21 pairs where neither genome has MPME1 (negative controls), 420 cross-type pairs where exactly one genome contains MPME1, and 1,770 pairs where both genomes have MPME1. MPME1 was extracted from the BPs genome (GenBank EU568876) and MPME2 from Halo (GenBank DQ398042), both originally characterized by Sampson et al. (Sampson et al., 2009); we use only MPME1 in our suffix set ℳ_*T*_ to test target specificity.

### Implementation

For full genome comparison (∼41 kb sequences), the *O*(*pq*) dynamic programming requires ∼1.7 *×* 10^9^ cell evaluations per pair. We implemented an optimized algorithm using Numba JIT compilation that precomputes suffix positions before the DP phase, achieving ∼50*×* speedup over pure Python (∼34 seconds vs. ∼30 minutes per mlev computation). Pairwise computations were parallelized across the NYU Greene HPC cluster (16 GB memory per job) using SLURM job arrays. Each of the 2,211 pairs was computed independently with both MPME1 and MPME2 motifs, sweeping the discount parameter *m* over {0.0, 0.1, 0.2, …, 1.0}. Total computation required approximately 500 core-hours.

## Results

We quantify the effect of mlev using the *discount percentage*: for each pair (*a, b*), discount = (lev(*a, b*) − mlev(*a, b*))*/* lev(*a, b*) *×* 100%, representing the fraction of edit distance attributable to discounted mobile element operations. Table 2 shows representative pairwise results at *m* = 0.5 and *m* = 1.0.

**Table 1.** Cluster G1 mycobacteriophage genome sizes and inferred MPME status. Phages with genomes ∼ 41,077–41,456 bp lack mobile elements; phages with longer genomes (∼ 41,650–42,400 bp) are inferred to contain MPME1 or a related element. *Halo carries MPME2 rather than MPME1; 14 phages with longer genomes lack both elements (see text).

| Phage | Size (bp) | Inferred MPME |
| --- | --- | --- |
| Angel | 41,441 | No |
| CLED96 | 41,456 | No |
| Cherrybomb426 | 41,456 | No |
| GoldenAsh | 41,441 | No |
| Schiebel | 41,441 | No |
| Taheera | 41,077 | No |
| Terror | 41,077 | No |
| *60 phages with longer genomes (representative examples):* |  |  |
| BPs | 41,901 | Yes |
| Hope | 41,901 | Yes |
| Zombie | 41,901 | Yes |
| BruceB | 41,901 | Yes |
| Halo | 42,289 | Yes* |

**Table 2.** Representative pairwise distances. “Neither” = both lack MPME1 (negative controls); “Cross (canonical)” = one has canonical MPME1, one does not; “Cross (MPME2)” = one has MPME2 instead of MPME1; “Both” = both contain MPME1. Discount is computed at *m* = 1.0. The Halo pair demonstrates that mlev correctly distinguishes MPME1 from the related element MPME2 (see text).

| Pair | Category | lev | mlev<sub>$m=0.5$</sub> | mlev<sub>$m=1$</sub> | Disc. |
| --- | --- | --- | --- | --- | --- |
| Angel vs Schiebel | Neither | 1 | 1.0 | 1.0 | 0.0% |
| Angel vs GoldenAsh | Neither | 67 | 67.0 | 67.0 | 0.0% |
| CLED96 vs Terror | Neither | 6061 | 5951.5 | 5842.0 | 3.6% |
| Angel vs BPs | Cross (canonical) | 618 | 400.5 | 183.0 | 70.4% |
| Angel vs Hope | Cross (canonical) | 619 | 401.5 | 184.0 | 70.3% |
| BPs vs Schiebel | Cross (canonical) | 617 | 399.5 | 182.0 | 70.5% |
| Angel vs Halo | Cross (MPME2) | 1582 | 1523.0 | 1464.0 | 7.5% |
| BPs vs BruceB | Both | 50 | 50.0 | 50.0 | 0.0% |
| BPs vs Hope | Both | 581 | 457.0 | 23.0 | 96.0% |
| BPs vs Phish | Both | 822 | 459.5 | 26.0 | 96.8% |
| Frosty24 vs Hope | Both | 702 | 618.5 | 343.0 | 51.1% |

Table 3 summarizes the mean discount by category for cross-type comparisons (reference phages vs. mobile element carriers, 420 pairs).

**Table 3.** Bidirectional validation of mlev using MPME1 (434 bp motif) and MPME2 (440 bp) as target motifs. Cross-type comparisons between the 7 original no-element reference phages and mobile element carriers reveal clear discrimination: MPME1 carriers (35 phages) show high discount with MPME1 motif but low with MPME2; MPME2 carriers (11 phages) show the reverse pattern. The “No-element vs Neither” row (98 pairs = 7 × 14) compares the 7 references against 14 phages newly identified by mlev as lacking both elements. This bidirectional approach confirms mobile element identity through sequence matching rather than genome size.

| Comparison Type | Pairs | Mean Discount ($m = 1.0$) | |
| --- | --- | --- | --- |
|  |  | MPME1 motif | MPME2 motif |
| No-element vs MPME1 carriers | 245 | $48.8\% \pm 34.0\%$ | $3.5\% \pm 1.2\%$ |
| No-element vs MPME2 carriers | 77 | $9.0\% \pm 3.7\%$ | $51.7\% \pm 34.7\%$ |
| No-element vs Neither | 98 | $8.0\% \pm 6.9\%$ | $2.8\% \pm 1.1\%$ |

Standard Levenshtein treats Angel–BPs, Angel–Halo, and BPs– Hope as roughly equidistant (lev ≈ 600), but mlev reveals fundamentally different biology: mobile element insertion (70% discount), a distinct element MPME2 (7% discount), and positional polymorphism (96% discount), respectively. Figure 1 confirms that the discount scales linearly with *m*, as predicted by the cost function, with all categories converging to zero at *m* = 0 (standard Levenshtein). The parameter *m* provides tunability: lower values weight mobile element events more heavily relative to point mutations. We report *m* = 1 throughout, which treats a full block insertion as a single edit operation and provides the clearest biological interpretation.

**Figure 1.**
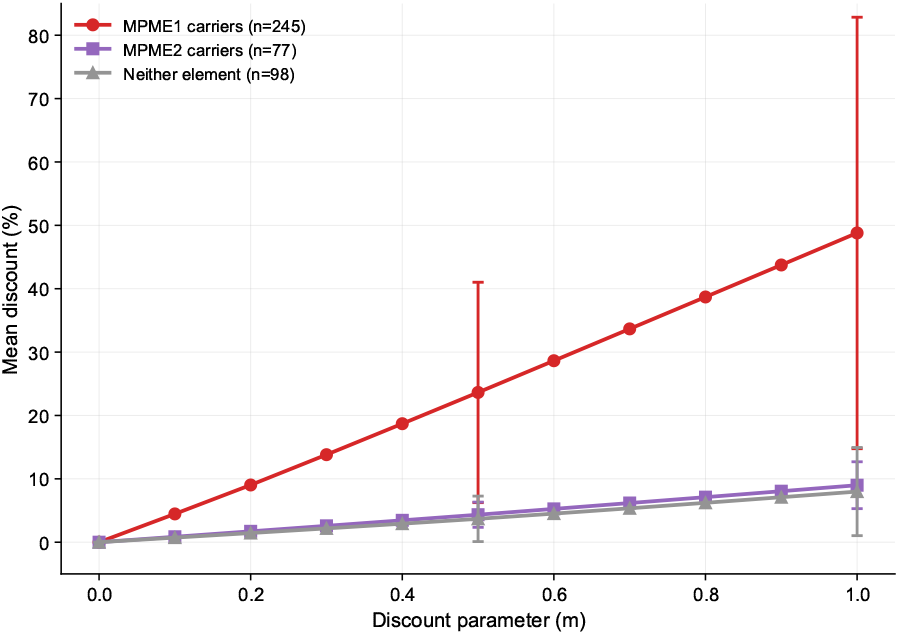
Mean discount percentage as a function of the discount parameter *m* for cross-type comparisons (reference phages vs. mobile element carriers). All three categories show perfect linear scaling through the origin, confirming theoretical predictions: at *m* = 0 (standard Levenshtein), discount is zero; at *m* = 1 (full discounting), MPME1 carriers reach ∼49% mean discount. The large error bars for MPME1 carriers reflect the bimodal distribution shown in Figure 3 (canonical vs. variant carriers). MPME2 carriers and phages with neither element follow nearly identical trajectories (∼8–9% at *m* = 1), demonstrating that the MPME1 motif is specific and does not cross-react— but also that these two groups cannot be distinguished by MPME1 analysis alone.

Figure 2 provides an overview of all 2,211 pairwise comparisons. The block structure immediately reveals the discriminative power of mlev: bright yellow blocks (high discount) appear wherever MPME1 carriers are compared to reference phages, while MPME2 and Neither blocks remain uniformly dark. Most striking is the checkerboard pattern within the MPME1 *×* MPME1 block: dark cells represent pairs where both phages carry MPME1 at the same genomic position (yielding zero net insertion difference), while yellow cells indicate different insertion positions—positional polymorphism that standard edit distance cannot detect.

**Figure 2.**
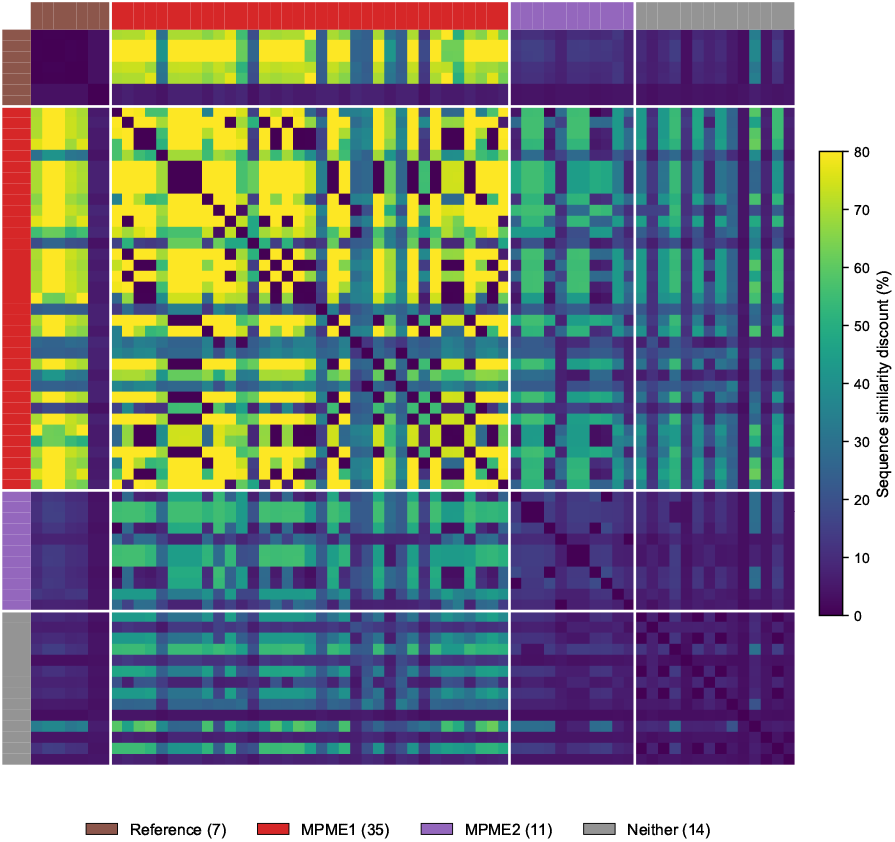
Heatmap of pairwise discount percentages at *m* = 1.0 for all 67 Cluster G1 phages (MPME1 motif). Phages are grouped by mobile element status: reference/no element (brown, 7 phages), MPME1 carriers (red, 35 phages), MPME2 carriers (violet, 11 phages), and neither element (gray, 14 phages). White lines delineate category boundaries. The block structure reveals clear discrimination: Reference × MPME1 blocks show high discount (yellow, ∼70%), while Reference × Reference, MPME2, and Neither blocks are uniformly dark (*<*10%). The MPME1 × MPME1 block (center) displays a striking checkerboard pattern: dark cells indicate pairs where both phages carry MPME1 at the *same* genomic position (no net insertion difference), while yellow cells indicate *different* positions—positional polymorphism detectable only by mlev. The diagonal is uniformly dark (self-comparisons yield zero discount).

### Validation and key findings

#### Method validation

Negative controls (21 pairs where neither genome contains MPME1) show minimal discount: mean 1.9% at *m* = 1, likely reflecting chance suffix matches with the 434-suffix motif set; this serves as the noise floor. Cross-type pairs with canonical MPME1 show the expected discount: at *m* = 1, a 434 bp block edit costs *c*(434) = 1 instead of 434 single-character operations, yielding an expected absolute discount of 433 edits. For a typical cross-type pair with lev ≈ 618, this corresponds to 433*/*618 ≈ 70%; observed values for canonical carriers (e.g., Angel vs BPs at 70.4%) closely match this prediction.

Figure 3 reveals that the MPME1 carrier distribution is *bimodal*, not normal: canonical carriers cluster at 65–100% while variant carriers produce a second peak at 0–20%. The bimodal structure is quantitatively evident: 261 pairs (62%) fall in the low mode (*<*25% discount) while 135 pairs (32%) fall in the high mode (*>*50% discount), with only 24 pairs (6%) in between. This bimodality explains the category mean (48.8%) and high standard deviation (*±*34%)—the mean does not represent a typical value but rather falls between two distinct subpopulations. Importantly, MPME2 carriers and phages with neither element show nearly identical unimodal distributions (means 9.0% and 8.0%), confirming that the MPME1 motif does not cross-react with other sequences but also demonstrating that MPME1-based analysis alone cannot distinguish MPME2 carriers from phages lacking mobile elements entirely.

**Figure 3.**
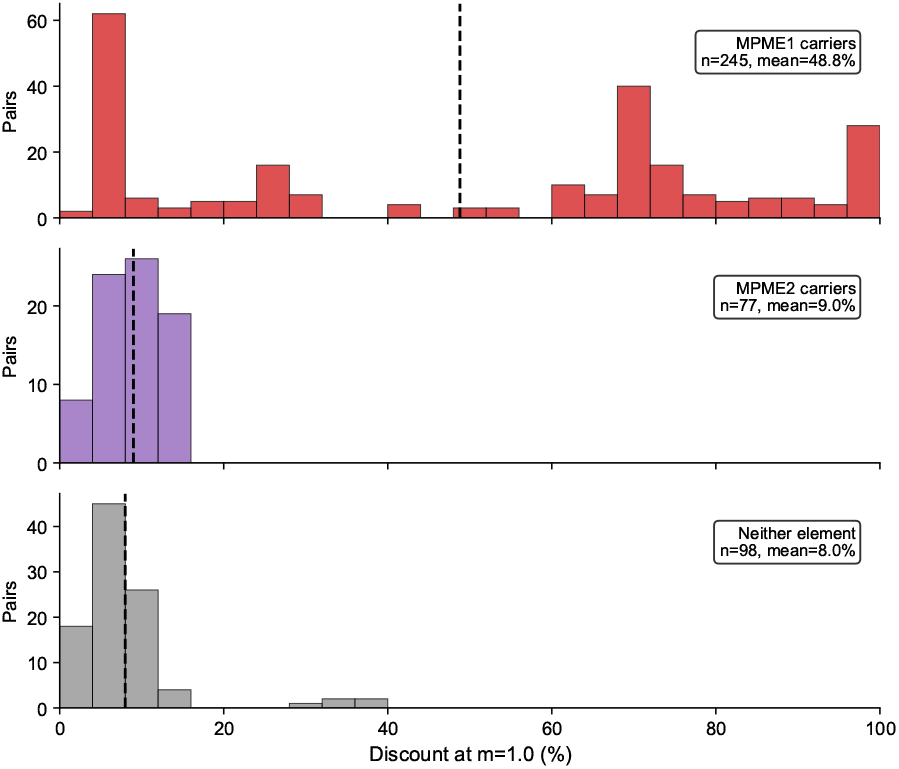
Distribution of discounts in cross-type comparisons (reference phage vs. mobile element carrier) using MPME1 motif. MPME1 carriers (red, *n* = 245) show a striking **bimodal distribution**: canonical carriers cluster at 65–100% discount while variant carriers and positional polymorphisms produce a second peak at 0–20%. This bimodality explains the high standard deviation (*±*34%) and why the mean (48.8%) does not represent a typical value. MPME2 carriers (violet, *n* = 77) and phages with neither element (gray, *n* = 98) both show unimodal distributions clustered at low values (5– 15%), confirming no cross-reactivity with the MPME1 motif; these groups are indistinguishable by MPME1 analysis alone, motivating the bidirectional validation in Figure 4.

#### Discovery: Bidirectional MPME1/MPME2 discrimination

To validate our discrimination of MPME1 from MPME2, we performed a second analysis using the MPME2 element (440 bp, extracted from Halo genome DQ398042) as the target motif. Figure 4 shows each non-reference phage plotted by its mean discount (averaged across all 7 reference phages) with the MPME1 motif (x-axis) versus the MPME2 motif (y-axis). The three categories separate cleanly into distinct quadrants with no overlap across the 20% decision boundaries. MPME1 carriers show significantly higher discount with the MPME1 motif than MPME2 carriers (Mann-Whitney *U* = 385, *p* < 4 × 10^−7^), while MPME2 carriers show significantly higher discount with the MPME2 motif (Mann-Whitney *U* = 385, *p* < 4 × 10^−7^). A Kruskal-Wallis test confirms significant differences among all three groups (*H* = 42.4, *p* < 7 × 10^−10^). Statistical tests were performed using SciPy (Virtanen et al., 2020).

**Figure 4.**
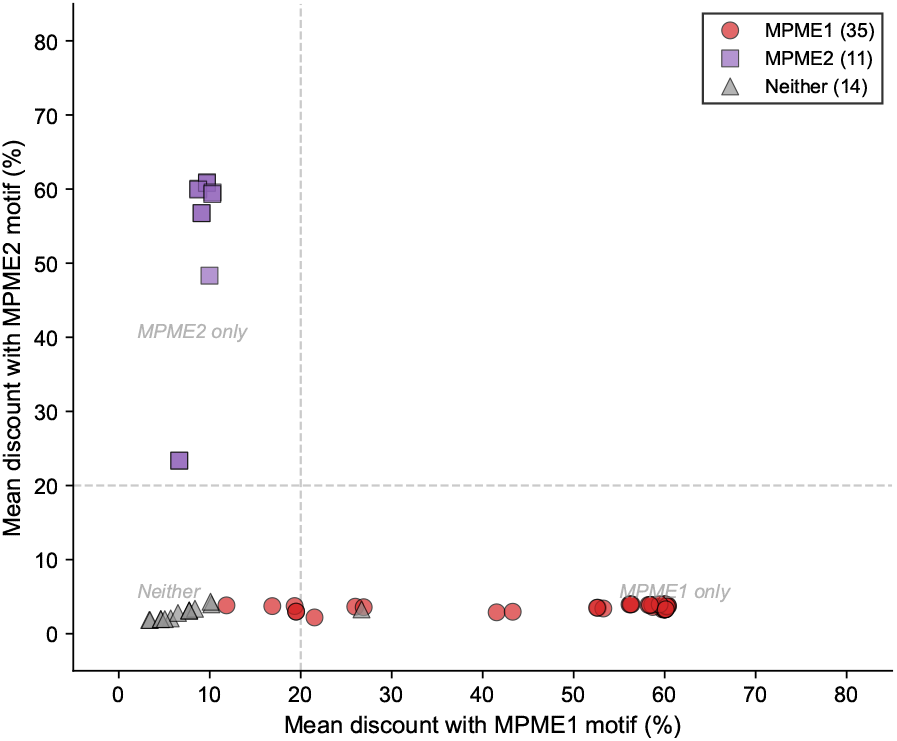
Bidirectional validation scatter plot. Each point represents one of 60 non-reference phages; coordinates are mean discounts averaged across comparisons with all 7 reference phages. X-axis: discount with MPME1 motif; Y-axis: discount with MPME2 motif. The three mobile element categories separate into distinct quadrants: MPME1 carriers (red circles, *n* = 35) cluster in the lower-right (high MPME1 discount, low MPME2), MPME2 carriers (violet squares, *n* = 11) in the upper-left, and phages with neither element (gray triangles, *n* = 14) in the lower-left corner. Dashed lines at 20% indicate approximate decision boundaries.

MPME1 carriers (red circles) cluster in the lower-right quadrant with high MPME1 discount (20–65%) but uniformly low MPME2 discount (∼3–5%). The wide spread along the x-axis reflects sequence variation: canonical carriers cluster near 60%, while eight phages with positional polymorphism or sequence variants (e.g., Frosty24 with a single SNP) show intermediate values. MPME2 carriers (violet squares) show the inverse pattern in the upper-left quadrant: low MPME1 discount (∼7–12%) but high MPME2 discount (23–62%). The outlier at ∼23% is Halo, from which the MPME2 motif was extracted; this lower self-discount likely reflects sequence divergence in the specific Halo isolate used. This orthogonality—each motif produces high discount only for its corresponding element—confirms that mlev discriminates between distinct mobile elements rather than detecting a generic insertion signal.

The 14 phages in the lower-left quadrant (gray triangles) show low discount with *both* motifs (*<*10% on both axes), forming a tight cluster near the origin. These phages—Aroostook, Barkley26, Chance64, Gideon, Grizzly, Hotshotbaby7, Jolene, Jonghyun, Liefie, Paito, Phloodle, Rabbs, Ruriko, and Sneeze—show low motif-specific discount consistent with absence of both MPME1 and MPME2 sequences (or highly divergent copies), despite having genome sizes larger than the no-element references. Whether they contain a distinct element, unrelated insertions, or other sequence differences remains to be determined. The final classification based on mlev discount patterns: 35 MPME1 carriers, 11 MPME2 carriers, 21 putatively lacking both elements (7 original references plus 14 identified by mlev).

#### Additional capabilities

When both genomes contain MPME1 at the same position, we observe zero discount; when at different positions (e.g., BPs vs Phish at 96.8%), mlev detects the deletion plus reinsertion— positional polymorphism invisible to standard edit distance. The phage Frosty24 carries an MPME1 variant with a single SNP at position 368, which breaks suffix matching and produces intermediate discounts (∼19% instead of ∼70%), demonstrating that mlev can detect single nucleotide differences within mobile elements.

#### Classification performance

To quantify the advantage of motif-specific discounting, we evaluated classification performance for detecting MPME1 carriers using receiver operating characteristic (ROC) analysis (Fawcett, 2006). To ensure statistical independence, we computed per-genome mean discounts (averaged across all 7 reference comparisons), yielding 60 independent observations rather than 420 non-independent pairs. We compared mlev discount to three standard pairwise distances: standard Levenshtein distance, affine gap alignment score, and 21-mer Jaccard distance. Figure 5 shows the resulting ROC curves. The mlev discount achieves AUC = 0.99, substantially outperforming Levenshtein (AUC = 0.78), affine gap (AUC = 0.73), and 21-mer Jaccard distance (AUC = 0.71). At the 20% discount threshold used throughout this paper, mlev achieves 96.0% specificity and 85.7% sensitivity; the imperfect sensitivity reflects sequence variants in some MPME1 carriers (e.g., Frosty24). This comparison demonstrates that motif-specific discounting captures biological signal—the presence of a specific mobile element—that bulk similarity measures cannot detect.

**Figure 5.**
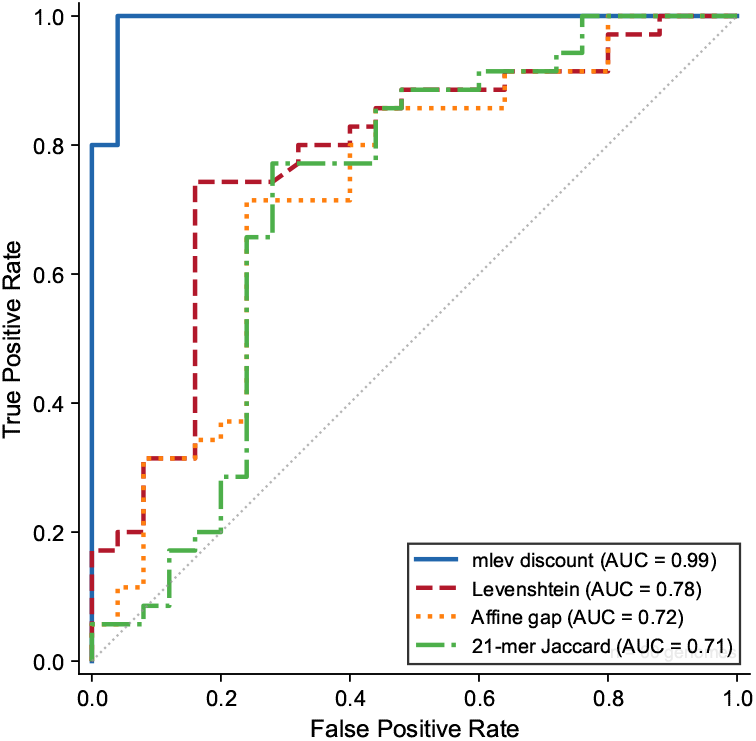
ROC curves for MPME1 carrier classification using per-genome mean discounts (*n* = 60 non-reference phages). Each phage’s score is its mean discount averaged across all 7 reference comparisons, ensuring statistical independence. The mlev discount (solid blue, AUC = 0.99) substantially outperforms standard Levenshtein distance (dashed red, AUC = 0.78), affine gap alignment (dotted orange, AUC = 0.73), and 21-mer Jaccard distance (dash-dot green, AUC = 0.71). The diagonal dotted line represents random classification (AUC = 0.50).

#### Independent sequence validation

To validate the mlev classifications independently, we performed direct sequence searches for MPME1 and MPME2 motifs in each genome. Of 35 phages classified as MPME1 carriers by mlev, 31 (89%) contain ≥90% of the MPME1 motif sequence; the four exceptions (Camri, Frosty24, Phreak, Renaissance) have sequence variants or positional differences that reduce exact matching but are still detected by mlev’s suffix-based approach. All 11 MPME2 carriers contain ≥90% of the MPME2 motif. All 7 reference phages and 13/14 “neither” phages show *<*50% match to both motifs; the exception (Phloodle, 62% MPME1 match) may represent a highly divergent or truncated element. Overall agreement between mlev classification and direct sequence search is 92.5% (62/67 phages).

## Discussion

### Comparison with existing methods

Table 4 summarizes the capabilities of mlev relative to existing approaches. Standard edit distances (Levenshtein, affine gap) produce pairwise distances but cannot distinguish mobile element insertions from random indels. Mobile element detection tools (MELT, RetroSeq) identify insertions but require short-read sequencing data and do not produce pairwise distances. Block edit distances allow substring operations but are NP-hard in general and not motif-specific. The mlev framework uniquely combines four capabilities: (1) pairwise distance, (2) motif-specific discounting, (3) positional polymorphism detection, and (4) polynomial-time computation from assembled sequences. The quantitative advantage is substantial: in ROC analysis for MPME1 carrier detection (Figure 5), mlev achieves AUC = 0.99 compared to 0.78 for standard Levenshtein, 0.73 for affine gap, and 0.71 for k-mer distance, demonstrating that motif-specific discounting captures biological signal that bulk similarity measures miss.

**Table 4.** Capability comparison of sequence comparison and mobile element detection methods. “Pairwise distance” = produces quantitative pairwise distances; “Motif-specific” = discounts only user-specified element sequences; “Positional polymorphism” = detects same element at different genomic locations; “Poly-time” = polynomial time complexity; “Input type” = required input data. Only mlev combines all four capabilities from assembled sequences.

| Method | Pairwise distance | Motif-specific | Positional polymorphism | Poly-time | Input type |
| --- | --- | --- | --- | --- | --- |
| Levenshtein | ✓ | — | — | ✓ | Sequence |
| Affine gap (Gotoh, 1982) | ✓ | — | — | ✓ | Sequence |
| Block edit (Lopresti and Tomkins, 1997) | ✓ | — | — | — | Sequence |
| MELT (Gardner et al., 2017) | — | ✓ | — | ✓ | Short reads |
| RetroSeq (Keane et al., 2013) | — | ✓ | — | ✓ | Short reads |
| mlev | ✓ | ✓ | ✓ | ✓ | Sequence |

### Limitations

The current implementation has several limitations. First, the *O*(*pq*) complexity restricts practical application to genomes under ∼50 kb; bacterial chromosomes (∼5 Mb) would require approximate methods. Second, mlev detects only exact suffix matches to the motif set; SNPs within mobile elements (as seen with Frosty24) reduce the discount, and highly divergent element copies may escape detection entirely. Third, the suffix-based approach cannot distinguish between a true mobile element insertion and an unrelated sequence that happens to match the motif suffixes by chance—though our negative controls suggest this occurs rarely (*<*2% discount in pairs lacking theelement). Finally, the asymmetry introduced by the retrotransposon model (discounted insertion but not deletion) means mlev(*a, b*) ≠ mlev(*b, a*) in general; all results in this paper use the symmetric version (see Section 2.2).

### Biological implications

The bimodal distribution of discounts among MPME1 carriers (Figure 3) reveals unexpected heterogeneity: rather than a single canonical element, the MPME1 family includes positional variants (different insertion sites), sequence variants (SNPs within the element), and possibly truncated copies. This heterogeneity would be invisible to standard edit distance, which treats all 35 MPME1 carriers as roughly equidistant from the references. The discovery of 14 phages carrying neither MPME1 nor MPME2 despite having expanded genomes (*>*41.5 kb) raises intriguing questions: do these phages harbor a third, uncharacterized mobile element? Or do their size differences reflect accumulated small indels unrelated to mobile element activity?

### Future directions

Natural extensions include: incorporating multiple motif variants in the suffix sets to track several mobile element families simultaneously; applying approximate methods (suffix-tree acceleration (Ukkonen, 1995) or seed-and-extend heuristics (Altschul et al., 1990)) to scale to larger genomes; and adapting mlev with appropriate motif sets for viral genome reconstruction, where recombination events create block-level mutations analogous to transposition. The framework could also be extended to weight different motif families differently, allowing simultaneous tracking of elements with different biological impacts.

## Conclusion

We formalized transposon and retrotransposon mechanisms as a mosaicism grammar and introduced a modified Levenshtein distance mlev with tunable block-event discounting. Validation on 67 mycobacteriophage genomes confirmed theoretical predictions: cross-type pairs with canonical MPME1 show the expected ∼70% discount, while negative controls remain near zero. Bidirectional analysis using both MPME1 and MPME2 as target motifs classifies 35 MPME1 carriers and 11 MPME2 carriers, and identifies 14 phages with low discount against both motifs, consistent with absence of both elements.

## Abbreviations

MPME: Mycobacteriophage Mobile Element
mlev: modified Levenshtein distance
DP: dynamic programming

## Data Availability

The mlev implementation is available at https://doi.org/10.5281/zenodo.18452982 (source code: https://github.com/kuhlman-lab-ucr/Genetic-Mosaicism). Mycobacteriophage genome sequences were obtained from PhagesDB (https://phagesdb.org). The MPME1 motif sequence (434 bp) was extracted from the BPs genome (GenBank accession EU568876, coordinates 39870–40303), and the MPME2 motif sequence (440 bp) was extracted from the Halo genome (GenBank accession DQ398042, coordinates 38875–39314). All pairwise distance results and analysis scripts are included in the repository.

## Author Contributions

H.K.S. developed the mathematical framework, implemented the algorithm, performed the computational analysis, and wrote the manuscript. T.E.K. supervised the research and revised the manuscript.

## Acknowledgments

We thank the graduate students in the Kuhlman Lab for helpful discussions: Aisa Anbir, Arya Naeini, and Shayan Nejad. Claude (Anthropic) was used for code development and editorial assistance.

